# Novel differential linear B-cell epitopes to identify Zika and dengue virus infections in patients

**DOI:** 10.1101/639542

**Authors:** Siti Naqiah Amrun, Wearn-Xin Yee, Farhana Abu Bakar, Bernett Lee, Yiu-Wing Kam, Fok-Moon Lum, Jeslin J.L. Tan, Vanessa W.X. Lim, Wanitda Watthanaworawit, Clare Ling, Francois Nosten, Laurent Renia, Yee-Sin Leo, Lisa F.P. Ng

**Affiliations:** Singapore Immunology Network, Agency for Science, Technology and Research (A*STAR), Singapore 138648, Singapore; Communicable Diseases Centre, Institute of Infectious Diseases and Epidemiology, Tan Tock Seng Hospital, Singapore 308433, Singapore; Shoklo Malaria Research Unit, Mahidol-Oxford Tropical Medicine Research Unit, Faculty of Tropical Medicine, Mahidol University, Mae Sot, Thailand; Centre for Tropical Medicine and Global Health, Nuffield Department of Medicine, University of Oxford, Oxford, UK; Lee Kong Chian School of Medicine, Nanyang Technological University, Singapore 636921, Singapore; Saw Swee Hock School of Public Health, National University of Singapore, Singapore 117549, Singapore; National Institute of Health Research, Health Protection Research Unit In Emerging and Zoonotic Infections, Liverpool, UK; Institute of Infection and Global Health, University of Liverpool, Liverpool L69 7BE, UK; Department of Biochemistry, Yong Loo Lin School of Medicine, National University of Singapore, Singapore 117597, Singapore

**Keywords:** Flavivirus, epitopes, patients, diagnostic

## Abstract

**Background:** Recent Zika virus (ZIKV) outbreaks challenged existing laboratory diagnostic standards, especially for serology-based methods. Due to the genetic and structural similarity of ZIKV with other flaviviruses, this results in cross-reactive antibodies which confounds serological interpretations.

**Methods:** Plasma from Singapore ZIKV patients was screened longitudinally for antibody responses and neutralizing capacities against ZIKV. Samples from healthy controls, ZIKV and DENV patients were further assessed using ZIKV and DENV peptides of precursor membrane (prM), envelope (E) or non-structural 1 (NS1) viral proteins in a peptide-based ELISA for epitope identification. Identified epitopes were re-validated and diagnostically evaluated using sera of patients with DENV, bacteria or unknown infections from Thailand.

**Results:** Long-lasting ZIKV-neutralizing antibodies were elicited during ZIKV infection. Thirteen potential linear B-cell epitopes were identified and of these, four common flavivirus, three ZIKV-specific, and one DENV-specific differential epitopes had more than 50% sensitivities and specificities. Notably, ZIKV-specific peptide 26 on domain I/II of E protein (amino acid residues 271-288) presented 80% sensitivity and 85.7% specificity. Importantly, the differential epitopes also showed significance in differentiating non-flavivirus patient samples.

**Conclusions:** Linear B-cell epitope candidates to differentiate ZIKV and DENV infections were identified, providing the first step towards the design of a much-needed serology-based assay.

## Introduction

Zika virus (ZIKV) outbreaks in French Polynesia and Brazil in 2013 and 2015 resulted in unexpected severe neurological and congenital complications [1–4], leading to a race to develop diagnostic and treatment strategies against the infection. Current ZIKV diagnosis, which relies heavily on molecular methods, poses several limitations because ZIKV patients display a short viremic phase with low viremia levels, and thus may escape detection, even in symptomatic patients [5,6]. Hence serology, as an alternative diagnostic approach, is very much needed to address these shortcomings. Unfortunately, this approach has been hampered due to the cross-reactive nature of the antibodies in ZIKV patients with other flaviviruses, such as dengue virus (DENV) [7–11], in which ZIKV shares high amino acid identity (55%) and structural homology with DENV [12–16]. Moreover, as both viruses are transmitted by the same mosquito vectors [17], they are often found in overlapping geographical areas [18,19]. Thus, there is a demand for a proper serology diagnostic tool that accurately differentiates the two infections.

Previous studies have shown the possibility of using ZIKV antigens to distinguish ZIKV infections from other flavivirus infections [11,20–22]. Although computational studies have predicted multiple differential epitopes, validation on patient samples however remains a challenge [23]. In this report, antibody and neutralizing responses by ZIKV patients from Singapore were characterized longitudinally. Common and differential linear B-cell epitopes recognized by antibodies from Singapore ZIKV and DENV patients were then identified. Importantly, the potential value of these identified epitopes in a diagnostic setting was further assessed using sera from patients from Thailand previously diagnosed with DENV, bacterial, and including those of unknown infections. This study aims to further the development of a serology-driven differential flavivirus diagnosis, particularly between ZIKV and DENV, allowing for accurate diagnosis that will improve patient management. The application can also be further expanded to study sero-prevalence and vaccine strategies.

## Methods

### Ethics statement

Written informed consent was obtained from participants in accordance with the tenets of the Declaration of Helsinki. Study protocols of Singapore ZIKV (2016-2018) and DENV (2010-2012) patient cohorts were approved by the SingHealth Centralized Institutional Review Board (CIRB Ref: 2016/2219) and National Healthcare Group (NHG) Domain Specific Review Board (DSRB-E-2009/432) respectively. Specimens from Singapore healthy donors (2010-2015) and patients from Thailand (2011-2013) were collected in accordance to study guidelines of approval numbers: NUS-IRB 09-256 and NUS-IRB 10-445; MUTM 2011-008-01, OXTREC 42-10 and TCAB-01-11 respectively.

### Study subjects and sample collection

#### Singapore ZIKV patients

Collection of specimens from subjects during the ZIKV outbreak in 2016 was previously described [24]. Briefly, 65 patients that were RT-PCR positive for ZIKV in whole blood or urine, and negative for DENV RT-PCR were enrolled [25]. Whole blood specimens were collected in EDTA-coated vacutainer tubes (Becton Dickinson) after peripheral venipuncture and were centrifuged at 12000 rpm for 10 min. Plasma was collected and heat-inactivated for 30 min at 56°C before storage at −80°C. Specimens were obtained over a period of six time points: (1) acute [2-7 days post-illness onset (pio)], (2) early convalescent (10-14 days pio), (3) late convalescent (1 month pio), (4) early recovery (3 months pio), (5) late recovery (5-6 months pio), and (6) full recovery (1 year pio) phases.

#### Singapore DENV patients

Twenty DENV patient serum samples (2010-2012) collected before the ZIKV outbreak were used in this study [26]. Patients were DENV PCR and/or NS1 positive upon hospital admission, and were a combination of the following: one unknown serotype, six DENV-1, seven DENV-2, three DENV-3, and three DENV-4 patients. Serum samples used were obtained at late convalescent phase (21-37 days pio).

#### Thailand patients

Archived serum samples from an undifferentiated fever study conducted at Shoklo Malaria Research Unit (SMRU) were used. Five DENV patients were confirmed by gold standard paired serology, and all but one was DENV PCR positive. Five bacteria-infected patients were diagnosed with leptospirosis, scrub typhus, murine typhus or *Streptococcus pneumonia*e infections, or a combination of above, and all were DENV PCR and DENV NS1, IgM and IgG RDT negative. Eight patients with unknown diagnoses were negative for the above pathogens by serology, blood culture and PCR. Convalescent serum samples used were collected at 14-20 days pio.

#### Viruses

ZIKV Polynesian isolate (H/PF/2013) was obtained from the European Virus Archive (EVA). DENV-3 was used as a reference DENV serotype because it is widespread in Southeast Asia [27–30], and was kindly provided by the National Public Health Laboratory (NPHL), Singapore. CHIKV SGP011 was isolated from a patient [31]. Viruses were propagated in VeroE6 cells (ATCC) and purified via ultracentrifugation [32] before being titered by standard plaque assays in VeroE6 cells [33,34].

#### Virion-based ELISA

Antibody titers were determined by a virion-based ELISA as previously described [18,32,34–36]. Briefly, purified virus was immobilized on 96-well maxisorp microtiter plates overnight (Nunc). Wells were blocked with 0.05% PBST [0.05% Tween-20 (Sigma-Aldrich) in PBS] containing 5% skim milk (Nacalai Tesque) at 37°C for 1.5 h. Heat-inactivated patient and pooled healthy control plasma samples at 1:200 to 1:8000 dilutions prepared in PBST with 2.5% milk were incubated at 37°C for 1 h. HRP-conjugated goat anti-human IgM or IgG (H+L) (Invitrogen) or mouse anti-human IgG1, IgG2, IgG3 and IgG4 (Thermo Fischer Scientific) antibodies were used for detection. Reactions were developed using TMB (3,3,5,5-tetramethyl benzidine) substrate (Sigma-Aldrich) and terminated with Stop reagent (Sigma-Aldrich), and absorbance was measured at 450 nm in a microplate autoreader (Tecan) [18,32,34–36]. ELISA readings were conducted in duplicates or triplicates.

#### Sero-neutralization

Neutralizing capacity of antibodies from ZIKV patients were determined via flow cytometry [37]. Briefly, pooled patient and healthy plasma samples at 1:1000 dilution were incubated with ZIKV or DENV-3 at MOI 10 for 2 h at 37°C with gentle agitation (350 rpm). Virus-antibody suspensions were then added in duplicates to HEK 293T cells (ATCC) at 37°C. After 2 h, media were removed and Dulbecco’s Modified Eagle Medium (DMEM; HyClone) with 10% fetal bovine serum (FBS; HyClone) were added. After 48 h, cells were harvested and stained as described [37], using ZIKV NS3 protein-specific rabbit polyclonal antibody [38] or DENV human monoclonal antibody 1B [34], and counter-stained with fluorophore-tagged goat anti-rabbit or anti-human IgG (H+L) (Life Technologies) respectively. Cells were acquired with MacsQuant Analyzer 10 (Miltenyi-Biotec). Flow cytometry results were analyzed with FlowJo (version 10.4.1, Tree Star Inc). Data of patient and pooled healthy neutralization assays were normalized using the respective untreated infections and calculated as a percentage of virus-only control infection.

### Epitopes determination

#### Linear peptide libraries

The sequences used for the design of biotinylated linear peptides of prM, E and NS1 proteins were derived from ZIKV Polynesian isolate (KJ776791) and consensus sequence of DENV-3 strains (KR296743, KF973487, EU081181, KF041254, JF808120, JF808121, KJ189293, KC762692, KC425219, KJ830751, KF973479, and AY099336) [32,34,36]. Peptides were generated as a ZIKV and DENV peptide-pair of corresponding sequences. Preliminary epitope screening was used with a library of peptides (Mimotopes) consisting of 18-mer overlapping sequences. Five peptides were combined to form one pooled peptide set. Screening and validation of patients were done with higher purity of peptides (≥90%, EMC microcollections GmbH) with lengths ranging from 11 to 22-mer (Supplementary Table 1). Peptides were dissolved in DMSO (Sigma-Aldrich) to obtain a stock concentration of 3.75 μg/μl.

#### Peptide-based ELISA

Epitope determination was performed via peptide-based ELISA as previously described [32,34,36]. Briefly, streptavidin-coated plates (Pierce) were blocked with 0.1% PBST (0.1% Tween-20 in PBS) containing 1% sodium caseinate (Sigma-Aldrich) and 1% bovine serum albumin (BSA; Sigma-Aldrich) overnight at 4°C, before addition of biotinylated peptides (1:1000 dilution in 0.1% PBST), followed by heat-inactivated pooled healthy control and patient plasma/serum samples (1:2000 dilution in 0.1% PBST). HRP-conjugated goat-anti human IgG (H+L) antibody (Invitrogen) prepared in 0.1% blocking buffer was used for detection of peptide-bound antibodies. TMB substrate and Stop reagent (Sigma-Aldrich) were used for development, prior to absorbance measurements at 450 nm (Tecan) [32,34,36]. All incubation steps were at room temperature for 1 h on a rotating shaker, and ELISA readings were conducted in duplicates.

#### Data analysis

OD values obtained from ZIKV and DENV peptide-based ELISA experiments were first normalized against mean OD values of pooled healthy donors. Patient samples were considered positive if the normalized response was more than 1.01. Subsequently, peptide binding capacity was calculated using the normalized values as [(ZIKV peptide response – DENV peptide response)/DENV peptide response]. Binding capacities with positive values denote the binding preference of the sample to ZIKV peptide, whereas negative values denote a binding preference to the corresponding DENV peptide. Difference in the mean peptide binding capacity of ZIKV patients and DENV patients of a peptide-pair (i.e. ZIKV and DENV peptides with complementary sequence) was calculated. Peptides with a relative difference of 0.1 or more are considered to be differential ZIKV (red) and DENV (blue) epitopes of interest, whereas peptides with a difference of 0.05 or less, and share amino acid similarity between the peptide-pair (Supplementary Table 1) are considered as common flavivirus epitopes (green).

#### Data visualization and statistical analysis

Heat-maps were generated using Multi Experiment Viewer (version 4.8, Microarray Software Suite TM4). For structural localization, data were retrieved from PDB 5IZ7 (ZIKV E) and 5K6K (ZIKV NS1). ZIKV prM and other DENV-3 proteins were simulated using Phyre [39]. For ZIKV prM, DENV-3 prM and E proteins, their structures were modeled based on PDB 4B03, and DENV-3 NS1 protein was modeled based on PDB 5K6K. All structures were visualized using PyMol (Schrodinger). Principal component analysis (PCA) was performed using the OD values of the anti-peptide IgG response by patients using prcomp function in R.

Statistics were done using GraphPad Prism (version 7.03). Mann-Whitney two-tailed tests with Bonferroni correction for multiple testing, or Kruskal-Wallis tests with Bonferroni correction for multiple testing, and post hoc tests using Dunn’s multiple comparison tests were used to derive any statistical significance. Correlation analysis was carried out using Spearman’s rank correlation. *P* values less than 0.05 are considered significant.

## Results

### ZIKV patients produce a robust and protective humoral response

Forty-five healthy donors were first screened for the presence of IgM and IgG against ZIKV, DENV and chikungunya virus (CHIKV), the three main arboviruses co-circulating in Singapore and several parts of Asia [18] using virion-based ELISA [18,32,34–36]. Twenty-two donors which had antibody levels lower than the assigned cut-off (mean + SD) in all three viruses (Supplementary Figures 1A-B) were used as the healthy control pool, and set as a baseline reference.

Anti-ZIKV IgM and IgG levels of ZIKV patients from the Singapore outbreak in 2016 [24,25,38] were longitudinally assessed using virion-based ELISA [18,32,34–36]. Majority of the patients showed a robust ZIKV-specific humoral response (Figures 1A-C, Supplementary Figure 1C). Anti-ZIKV IgM was detected as early as in the acute phase (2-7 days pio), and peaked at early convalescent (10-14 days pio), before decreasing during the recovery phases (3 months to 1 year pio) (Figures 1A and 1C, Supplementary Figure 1C). ZIKV-specific IgG titers peaked at early convalescent, persisted at high levels during late recovery, and were still detectable a year after infection (Figures 1B-C, Supplementary Figure 1C). These patients were also screened for the presence of DENV-specific antibodies and 80% of the patients were negative for anti-DENV IgM in samples taken at the acute phase (Supplementary Figures 1D and 1F). However, 75% of the patients were found to have anti-DENV IgG (Supplementary Figures 1E-F), suggesting that ZIKV IgG, but not IgM, cross-reacts with DENV.

**Figure 1.**
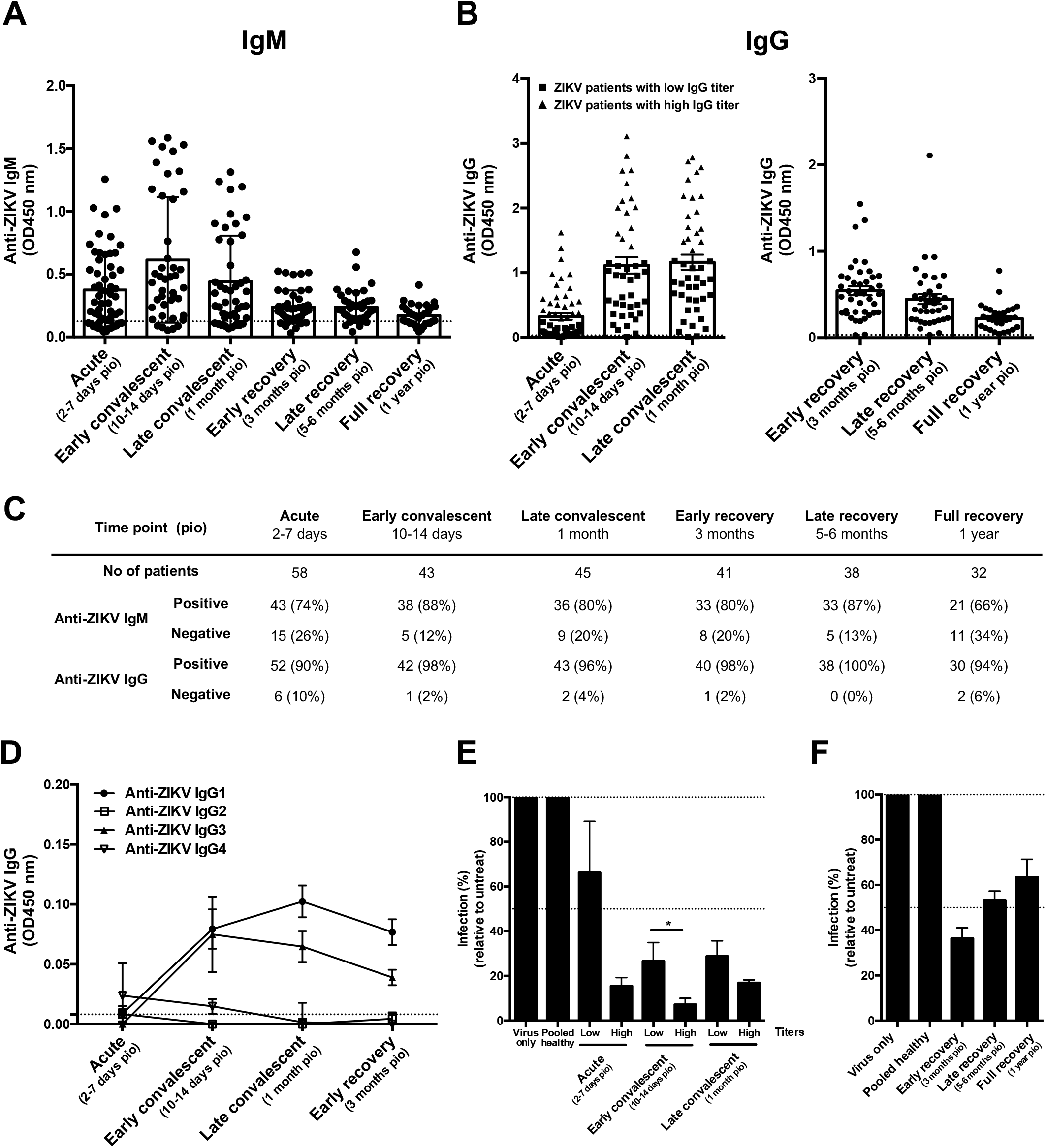
Antibody profiles of ZIKV patients of Singapore cohort in 2016 over time. (A-C) Total anti-ZIKV (A) IgM and (B) IgG antibody titers in patients’ plasma samples, at dilutions 1:200 and 1:2000 respectively, were determined by virion-based ELISA using purified ZIKV virions. Pooled plasma of healthy donors were used as negative control. Data are presented as mean ± SEM, with dotted line indicating mean of pooled healthy control. (C) Number and percentage of patients that are positive or negative for anti-ZIKV IgM and IgG at the respective time points. (D) IgG isotype titers in patients’ plasma samples were determined at 1:200 dilution in a ZIKV virion-based ELISA. Data are presented as mean ± SEM, with dotted line indicating mean of pooled healthy control. All ELISA readings were conducted in duplicates or triplicates. [Acute (n=58), early convalescent (n=43), late convalescent (n=45), early recovery (n=41), late recovery (n=38), full recovery (n=32)]. (E-F) *In vitro* neutralizing capacity of pooled ZIKV patients and pooled healthy control were tested at 1:1000 plasma dilution via flow cytometry. (E) Plasma samples were pooled according to levels of anti-ZIKV IgG titer [group of low titer patients are denoted as square symbol, while group of high titers are denoted as triangle symbol as shown in (B)] for acute [low (n=37), high (n=21)], early convalescent [low (n=29), high (n=14)], and late convalescent [low (n=28), high (n=17)] time points. (F) Plasma samples collected at the recovery phases were pooled together at the respective time points [early recovery (n=41), late recovery (n=38), full recovery (n=32)]. Results are expressed as percentage of control infection. Data presented as mean ± SD and representative of 2 independent experiments. Statistical analysis between low and high anti-ZIKV IgG titer groups was carried out using Mann-Whitney two-tailed test, with Bonferroni correction for multiple testing (\**p*<0.05).

IgG isotypes produced by ZIKV patients were then determined and highest titers of anti-ZIKV IgG1 and IgG3 subtypes were produced at early convalescent for IgG3, and late convalescent for IgG1 (Figure 1D). To determine if antibodies produced in these patients were protective against ZIKV, neutralization assays were carried out via flow cytometry. Efficient neutralization (71% to 93%) was observed in early and late convalescent stages (Figure 1E), whilst weak neutralization (37% to 47%) was seen in late and full recovery stages (Figure 1F). Neutralization capacity of ZIKV patients correlated with levels of anti-ZIKV IgG (Supplementary Figures 1C and 4A). Plasma from these patients only minimally neutralized DENV (Supplementary Figures 1G-H), indicating ZIKV-specificity.

### Identification of specific B-cell linear epitopes recognized by antibodies from ZIKV and DENV patients

Preliminary mapping of specific ZIKV and DENV epitopes was first performed in a peptide-based ELISA on the most antigenic flavivirus antigens: prM, E and NS1 [32,34,40], using pooled linear ZIKV and consensus DENV peptides. Plasma/serum samples of ZIKV and DENV patients [26] taken at the late convalescent phase were used as IgG levels were highest at this time point (Supplementary Figure 1C). Results specifically showed two common flavivirus (pools 1 and 21), six potential ZIKV-specific (pools 6, 10, 11, 16, 17 and 24) and one potential DENV-specific (pool 19) pools were identified within the ZIKV and DENV proteome (Supplementary Table 2, Supplementary Figure 2). Thereafter, new peptides selectively designed based on exposed residues and computational predictions were re-synthesized for subsequent experiments (Supplementary Table 1) [23].

Interestingly, results showed differences between pooled and individual peptides (Table 1, Figure 2). These differences could be due to interferences of the pooled peptides, while single peptides allowed for more enhanced specific binding. Nevertheless, six potential common flavivirus peptides were identified which displayed less than 0.05 relative difference in the binding capacity between ZIKV and DENV patients (peptides 7, 36, 38, 39, 46, 49) (Table 1, Figure 2, Supplementary Figure 3). These peptides were also selected based on the close similarity between the ZIKV and DENV peptide sequence (Supplementary Table 1). Additionally, three potential ZIKV-specific (peptides 3, 26 and 32), and four potential DENV-specific peptides (peptides 9, 17, 43 and 45) with a binding capacity difference of more than 0.1 were identified (Table 1, Figure 2, Supplementary Figure 3).

**Figure 2.**
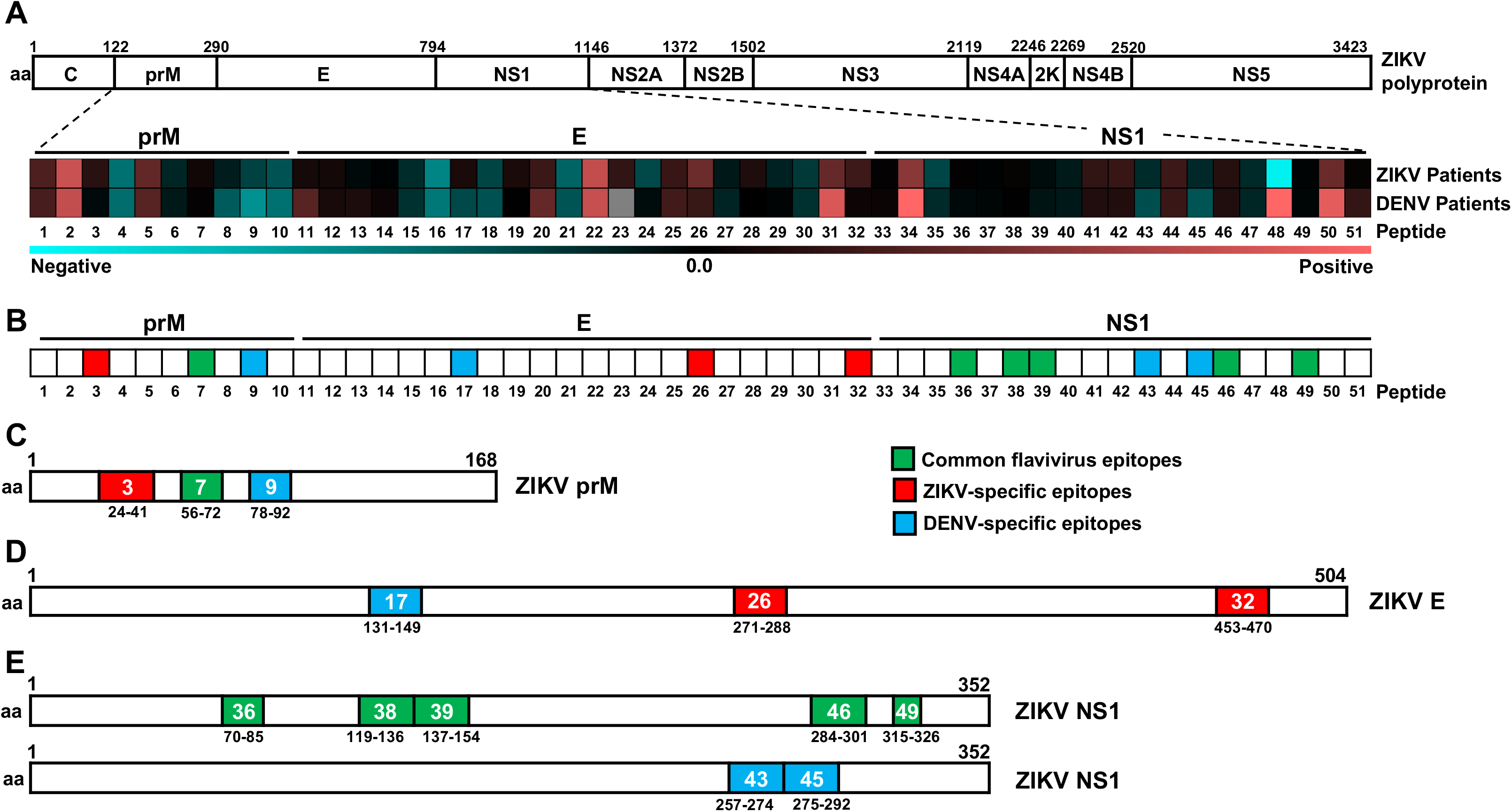
Mapping of common flavivirus, ZIKV-specific, and DENV-specific linear B cell epitopes using ZIKV and DENV patient samples. (A) Polyprotein of ZIKV H/PF/2013 (UniProtKB accession: A0A024B7W1). Plasma samples of ZIKV patients (n=30-44) and serum samples of DENV (n=20) patients at late convalescent phase were tested at 1:2000 dilution in a peptide-based ELISA in duplicates, using peptides that cover the precursor of membrane (prM: peptides 1-10), envelope (E; peptides 11-32) and non-structural 1 (NS1; peptides 33-51) proteins of ZIKV and DENV proteome. IgG response of patients were normalized to mean of pooled healthy control. Patients’ response to ZIKV and DENV peptide-pairs were compared and the mean binding capacity are presented in a heat-map. A value of 0 on the scale denotes patients showing equal binding response to a ZIKV and DENV peptide-pair, whereas values larger than 0 show preferential of patients to bind to ZIKV peptide. Values smaller than 0 show binding preference of patients to DENV peptide. (B) A schematic representation to denote common flavivirus (green), ZIKV-specific (red), and DENV-specific (blue) peptides across prM, E and NS1 based on heat-map analysis above. (C-E) Genome organization of ZIKV prM, E and NS1. Regions of amino acids corresponding to the identified linear B-cell epitopes in (C) prM, (D) E and (E) NS1 are shown, with green areas denoting common flavivirus, red denoting ZIKV-specific, and blue denoting DENV-specific epitopes. Numbers in colored boxes denote the peptide number, and the amino acid position in the respective proteome.

**Table 1.**
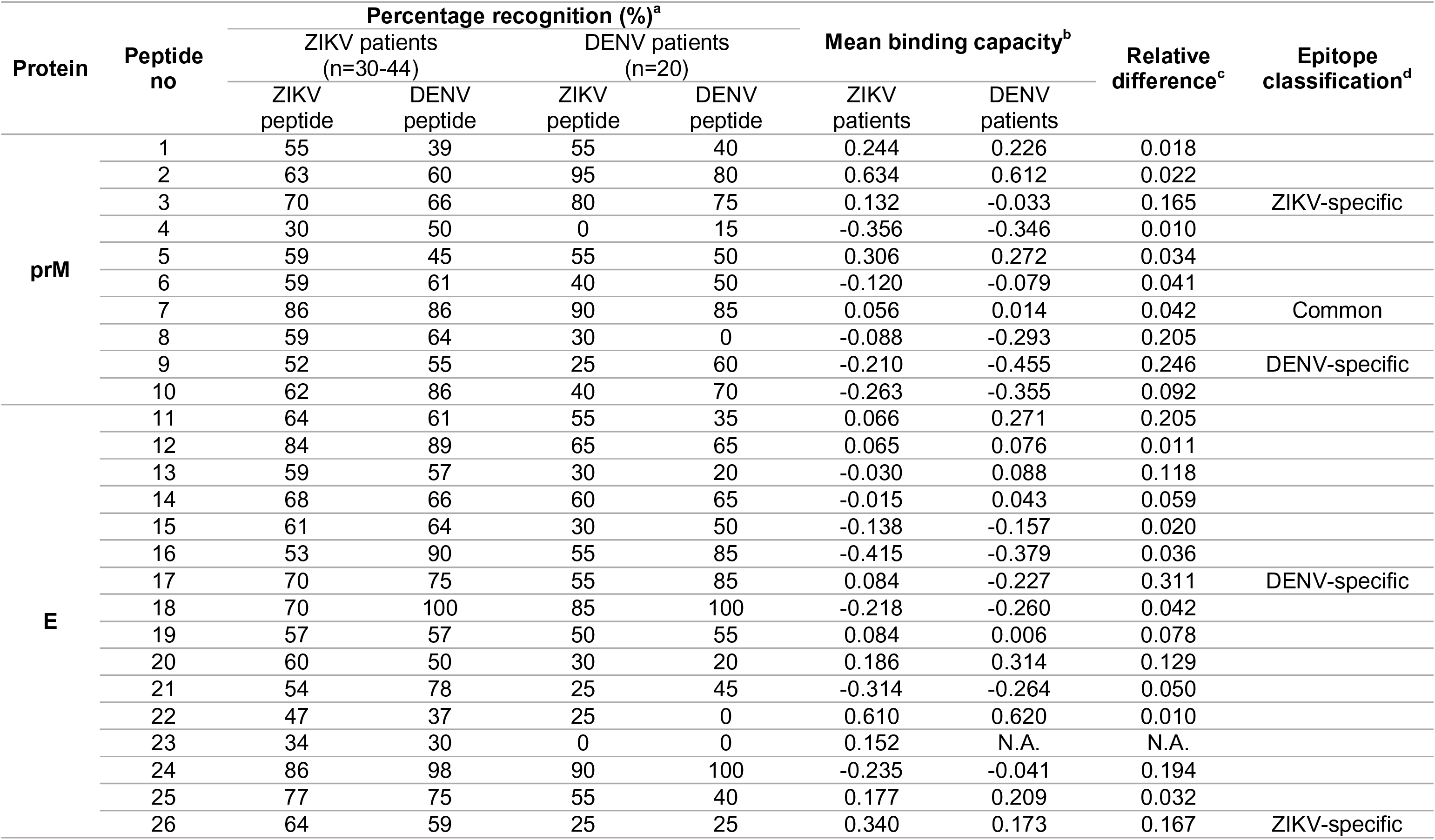
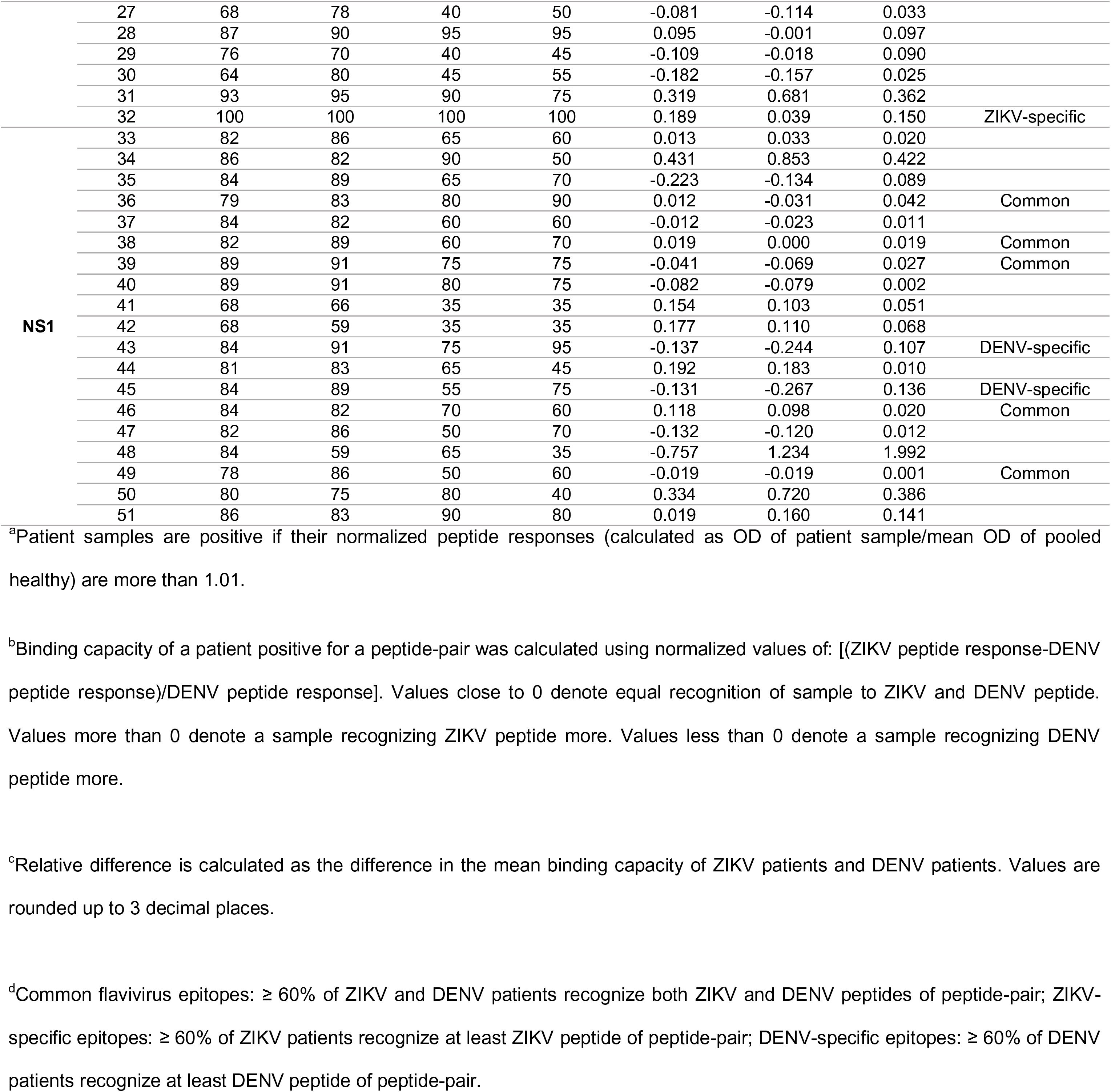
Singapore ZIKV and DENV patients’ response to ZIKV and DENV peptides

### Epitope recognition by ZIKV patients over time

In order to characterize the changes in epitope recognition by ZIKV patients over time, the common flavivirus (green) and ZIKV-specific peptides (red) were screened with plasma of ZIKV patients in acute, late convalescent, and full recovery phases. For the common flavivirus hits, more than 60% of the ZIKV patients were able to recognize the six peptide-pairs at late convalescent and beyond (Figure 3A). However, at the acute phase, only peptides 7, 36 and 38 were recognized by ZIKV patients (Figure 3A). In terms of binding capacity, there was equal binding between ZIKV and DENV peptide-pairs over time for peptides 7, 36, 38 and 49 (Figure 3B).

**Figure 3.**
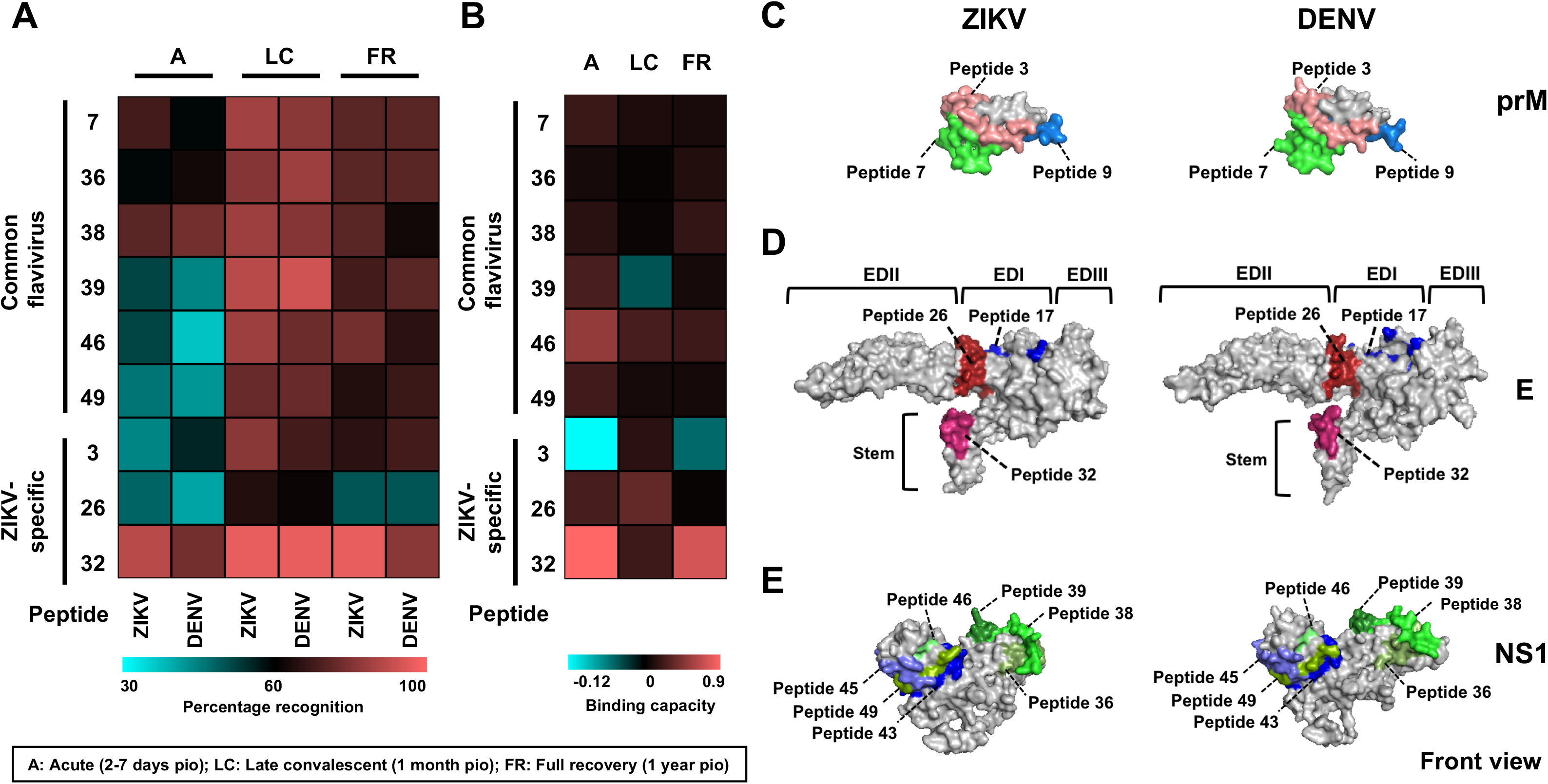
Characterization of the antibody profile kinetics of ZIKV patients on common flavivirus and ZIKV-specific linear B-cell epitopes, and localization of potential epitopes within the ZIKV and DENV proteome. (A-B) Plasma samples of ZIKV patients (n=27) at acute, late convalescent and full recovery phases were tested for IgG at 1:2000 dilution in duplicates using ZIKV and DENV peptides in a peptide-based ELISA. Pooled plasma of healthy donors was used as negative control and patients’ data were normalized to mean of pooled healthy control. (A) Percentage of ZIKV patients positively binding to ZIKV and DENV peptides, and (B) binding capacity of ZIKV patients positively binding to peptides were calculated and presented in a heat-map. (C-E) Schematic diagrams showing the localization of common flavivirus (denoted as shades of green), ZIKV-specific (denoted as shades of red), and DENV-specific (denoted as shades of blue) epitopes on (C) prM protein of ZIKV and DENV (PDB: 4B03), (D) E glycoprotein of ZIKV (PDB: 5IZ7) and DENV (PDB: 4B03), and (E) NS1 protein of ZIKV and DENV (PDB: 5K6K).

For ZIKV-specific epitopes, more than 60% of the ZIKV patient samples were able to recognize peptides 3 and 26 (Figure 3A), with positive peptide binding capacity (Figure 3B) at late convalescent phase. On the other hand, peptide 32 showed strong recognition by the patient samples (Figure 3A) as well as high binding capacity (Figure 3B) at various time points from acute to full recovery. The localization of all potential epitopes within the viral proteins are shown in Figures 3C-E.

### Evaluation of epitopes with patient cohorts

To assess the diagnostic performance of identified epitopes, the 13 peptides were screened using patient serum samples from a Thailand cohort that had DENV, bacteria, or unknown infections. Results of a randomized selection of Singapore ZIKV and DENV patients were also analyzed in parallel (Supplementary Table 3).

Interestingly, results showed a wide range of specificity and sensitivity for each peptide (Table 2, Figure 4A). ZIKV-specific peptide 26 (amino acid residues 271-288) on the E protein of domain I/II (EDI/II) had the best sensitivity and specificity profile (80% and 85.7% respectively) (Table 2, Figure 4A). Nevertheless, eight peptides (common flavivirus peptides 36, 38, 46, 49; ZIKV-specific peptides 3, 26, 32; and DENV-specific peptide 9) showed more than 50% sensitivity and specificity (Table 2, Figure 4A), and were selected for further evaluation. These peptides were used to “diagnose” the patients (Supplementary Table 4), and the performance of the peptide combination based on the epitope groupings were determined collectively (Table 2, Figure 4B). Although the common flavivirus (green) and DENV-specific (blue) groups demonstrated modest measurements, the ZIKV-specific (red) peptide mix showed a robust specificity of 96.4% (Table 2, Figure 4B). Furthermore, when the anti-peptide IgG response of patients was plotted in a principal component analysis (PCA), it was observed that patients of different diagnoses and cohorts formed separate clusters, and ZIKV patients stood out when compared to the healthy control (Figure 4C). To identify peptides with discriminating power, the binding capacity of positive peptides were calculated. The virus-specific ZIKV and DENV epitopes were significantly differential (Figure 4D). Peptide 32 (amino acid residues 453-470 on E protein) was the best performing ZIKV-specific epitope, and was able to distinguish Singapore ZIKV patients from bacteria and unknown infections from Thailand (Figures 4D-E). DENV-specific peptide 9 (amino acid residues 78-92 on prM) could be used to differentiate Singapore DENV patients from bacteria-infected patients from Thailand (Figure 4E). Overall, we have identified the best differential epitopes to differentiate between DENV and ZIKV patients.

**Figure 4.**
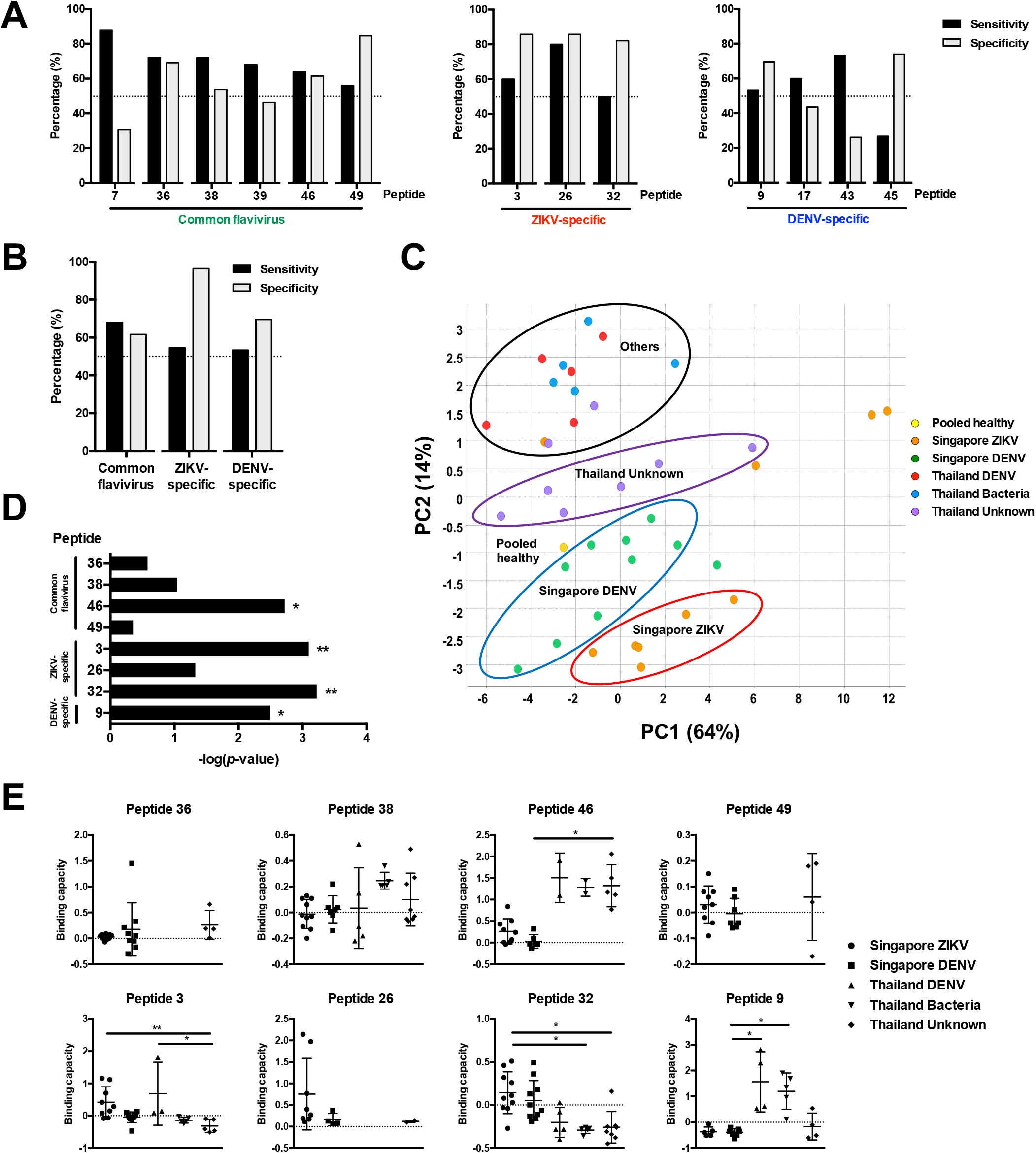
Preliminary diagnostic validation of identified linear B-cell epitopes with patient cohorts. Convalescent plasma samples of ZIKV (n=10) and serum samples of DENV (n=10) patients from Singapore, and DENV (n=5), bacteria (n=5) and unknown (n=8) patients from Thailand were tested in a peptide-based ELISA in duplicates at 1:2000 dilution. Pooled healthy plasma was used as a negative control. (A) Sensitivity and specificity were determined for individual peptides. (B) Sensitivity and specificity of peptide mix of selected epitopes were determined. (C) Principal component analysis (PCA) of pooled healthy and patients’ anti-IgG peptide response (OD values) were plotted in a graph with the percentage of variance indicated. (D-E) The peptide binding capacity of patients positively binding to peptides were calculated and statistically analyzed by using Kruskal-Wallis tests with Bonferroni correction for multiple testing. Post hoc tests were done using Dunn’s multiple comparison tests to determine (D) peptides with discriminating power, and (E) the peptide binding capacity distribution of patients. Data are presented as mean ± SD. (\**p*<0.05, \*\**p*<0.01).

**Table 2.**
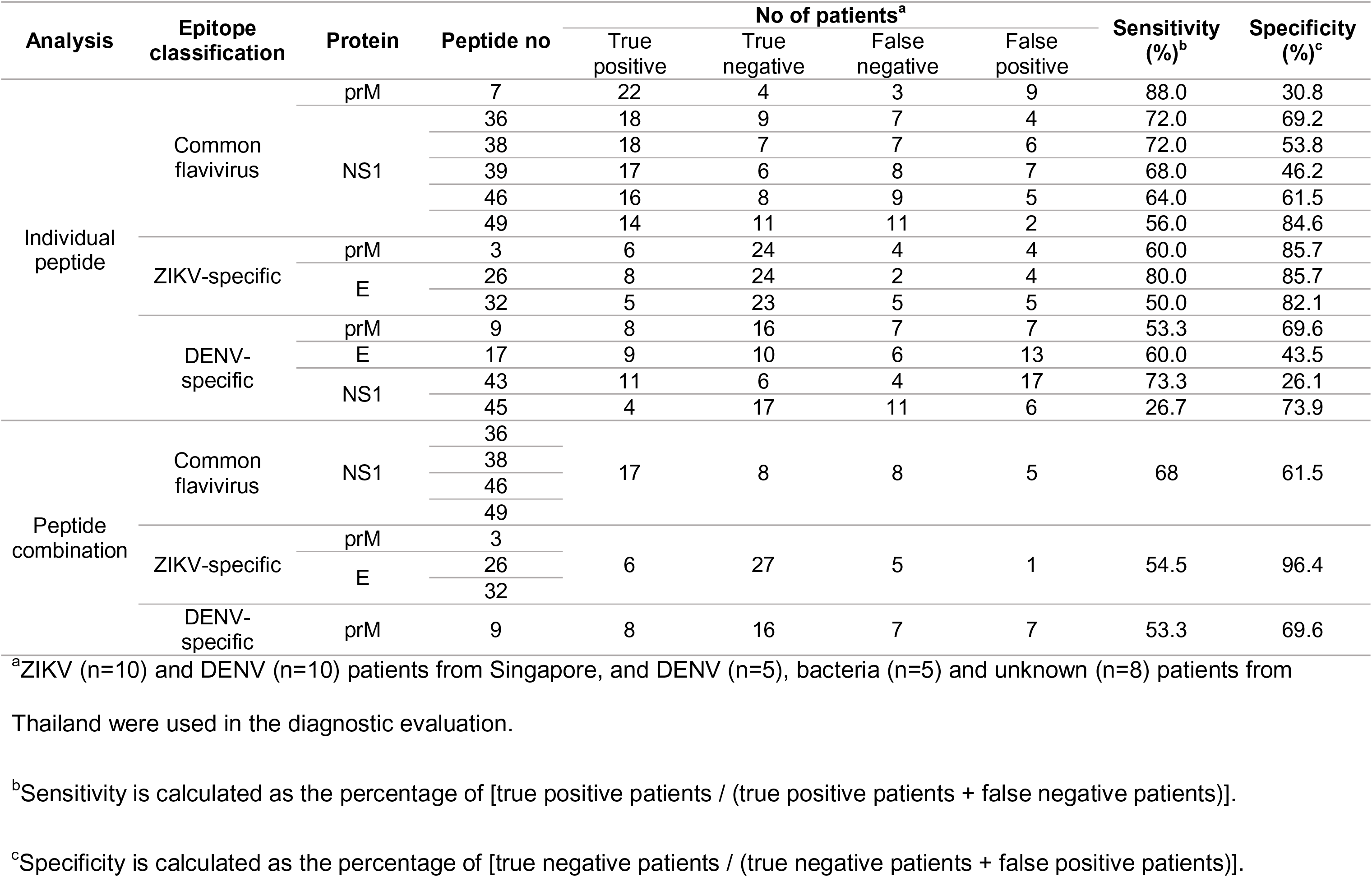
Diagnostic evaluation of linear B-cell epitopes

## Discussion

ZIKV patients were shown to produce high levels of ZIKV-specific IgG antibodies. Specifically, IgG1 and IgG3 were the subclasses induced following ZIKV infection, closely resembling DENV-infected patients [41]. Although patients from this cohort had detectable DENV IgG levels due to the high level of cross-reactivity among flaviviruses [7–10], DENV neutralization was significantly less efficient compared to ZIKV, indicating that the antibodies were ZIKV-specific (Figures 1E-F, Supplementary Figure 1G-H). This observation is also supported by another study, in which the profiles of ZIKV neutralizing antibodies of patients from Nicaragua, Sri Lanka and Thailand were not affected by previous DENV infection [42]. Nonetheless, it is imperative to consider the possible implications of virus-infection enhancement [43]. Moreover, none of the ZIKV patients in our study displayed severe symptoms to suggest occurrence of antibody-dependent enhancement (ADE) [24], and similar observations were also reported from Brazil [43,44].

While various reports have shown the specificity of the NS1 antigen to differentiate between ZIKV and DENV [11,20,21,45,46], majority of the common flavivirus peptides identified in this study are on the NS1 protein, possibly due to the conserved regions of NS1 amongst the flaviviruses [8,47]. For example, common flavivirus peptides 36 (amino acid residues 70-85), 38 (amino acid residues 119-136) and 49 (amino acid residues 315-326) were identified as ZIKV-specific in other patient cohorts from South America [45,46]. However, it remains to be seen if these peptides could be used to detect all flaviviruses such as yellow fever virus (YFV) and Japanese encephalitis virus (JEV).

Differential ZIKV and DENV epitopes identified were located across prM, E and NS1. Of interest, DENV-specific peptide 17 (amino acid residues 131-149) and ZIKV-specific peptide 26 are found on EDI and EDII of E glycoprotein, which share 35% and 51% amino acid identity between ZIKV and DENV respectively [8], whereas ZIKV-specific peptide 32 is located on the stem (Figure 3D). It would also be useful to assess the use of the identified peptides as a ZIKV vaccine target, particularly peptides 26 and 32. Interestingly, despite the similarity between the sequence of these ZIKV and DENV peptide-pairs (Supplementary Table 1), they were able to distinguish ZIKV and DENV patients. Moreover, ZIKV patients at different disease stages have different peptide recognition, and the current set-up could identify ZIKV infection at any point, independent of the patients’ level of ZIKV-specific antibodies (Supplementary Figure 4B-C). However, given that the identified epitopes were screened and validated using adult patient samples, it would be important to assess how these epitope profiles will perform in other patient cohorts, specifically ZIKV-infected pregnant women from Brazil [36].

Intriguingly, the Singapore DENV and Thailand DENV patients were not clustered together in the PCA (Figure 4B). Most of the Singapore DENV patients selected for validation had moderate to severe forms of plasma leakage, a clinical feature of severe manifestations of DENV infection [48], whereas DENV patients from Thailand displayed mild symptoms (unpublished data). The latter being “negative” in our assays could thus be due to differences in epitope recognition in different DENV disease states [40], and the different strain of viruses circulating in Singapore and Thailand. Nonetheless, further refinements are required to identify serotype-specific DENV epitopes.

Furthermore, comparing these results and computationally-predicted diagnostic peptide regions [23] revealed differences. Firstly, majority of the computationally predicted peptide regions were not ZIKV-specific. NS1 peptide 36, for example, was predicted to be differential [23], but was in fact a common flavivirus. However, peptides 26 and 32 on E protein, which were predicted to contain diagnostic epitopes [23], were indeed shown to be ZIKV-specific in this study. Thus, computational prediction remains useful to narrow down possible epitope candidates.

Overall, this study offers important valuable information on the human antibody response against ZIKV and insights into epitope cross-reactivity. Notably, several novel differential ZIKV and DENV epitopes with potential diagnostic efficacies have been identified on prM and E proteins. These results offer useful insights towards the development of diagnostics or vaccines.

## Acknowledgements

We thank Nicholas Q.R. Kng for assisting with the experiments, Kai-Er Eng for assisting with the manuscript, and Cheryl Y.P. Lee, Yi-Hao Chan, Guillaume Carissimo, Siew-Wai Fong and Jonathan Cox for processing of patient samples (SIgN); Vincent Pang, Linda Lee and research assistants from Tan Tock Seng Hospital; SMRU clinic and microbiology laboratory staff for patient recruitment, sample preparation and laboratory diagnosis, Mahidol-Oxford Tropical Medicine Research Unit (MORU) microbiology staff for supporting Rickettsia reference serology testing, and Armed Forces Research Institute of Medical Sciences (AFRIMS) for dengue reference serology testing; National Public Health Laboratory (NPHL) for providing DENV-3 virus; Prof Andres Merits for providing the ZIKV NS3 protein-specific rabbit polyclonal antibody; and study participants for their participation in the study.

