## Supplementary Figure 1 for "Novel differential linear B-cell epitopes to identify Zika and dengue virus infections in patients"

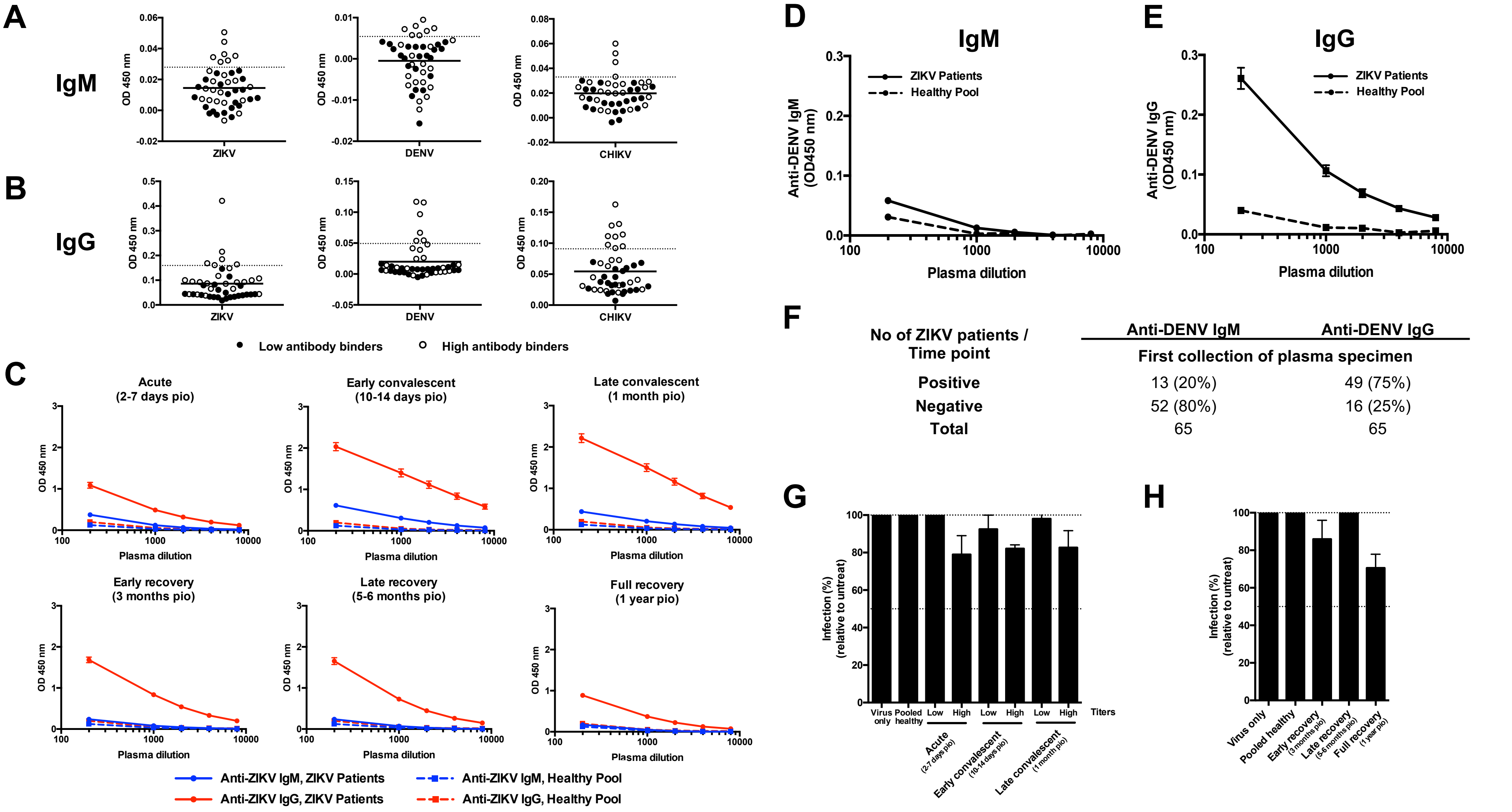


**Supplementary Figure 1. Antibody profiles of healthy controls and ZIKV patients of Singapore cohort in 2016.** Total (A) IgM and (B) IgG antibody titers of 45 healthy donors were determined by virion-based ELISA using purified ZIKV, DENV or CHIKV virions and plasma dilution of 1:2000. Data are presented as mean, with dotted line indicating the respective mean + SD values. Samples with OD values less than mean + SD for both IgM and IgG, and across all three viruses are highlighted in black (n=22), and were combined together to form the pooled healthy control for subsequent experiments. Samples with OD values more than the cut-off are denoted as clear symbols (n=23). (C) Total anti-ZIKV IgM and IgG of ZIKV patients were determined at plasma dilutions of 1:200, 1:1000, 1: 2000, 1:4000 and 1:8000 in virion-based ELISA using purified ZIKV virions at time points of acute (n=58), early convalescent (n=43), late convalescent (n=45), early recovery (n=41), late recovery (n=38), and full recovery (n=32). Pooled plasma of healthy donors was used as negative control. Data are presented as mean ± SEM. (D-E) Total anti-DENV (D) IgM and (E) IgG antibody titers in plasma samples of ZIKV patients at first collection time point were determined by virion-based ELISA using purified DENV virions in the same method as (C). Data are presented as mean ± SEM. All ELISA readings were in duplicates. (F) Number and percentage of patients that were positive or negative for anti-DENV IgM and IgG at the first collection of plasma specimen (1:200 and 1:2000 dilutions respectively). (G-H) *In vitro* neutralizing capacity of pooled ZIKV patients and pooled healthy control against DENV were tested at 1:1000 plasma dilution via flow cytometry. (G) Plasma samples were pooled according to levels of anti-ZIKV IgG titer (shown in Figure 1) for acute [low (n=37), high (n=21)], early convalescent [low (n=29), high (n=14)], and late convalescent [low (n=28), high (n=17)] time points. (H) Plasma samples collected at the recovery phases were pooled together at the respective time points [early recovery (n=41), late recovery (n=38), full recovery (n=32)]. Results are expressed as percentage of control infection. Data presented as mean ± SEM and representative of 2 independent experiments.

**Supp. Figure 1 Amrun_Yee *et al.*, 2019**
