## Supplementary Figure 2 for "Novel differential linear B-cell epitopes to identify Zika and dengue virus infections in patients"

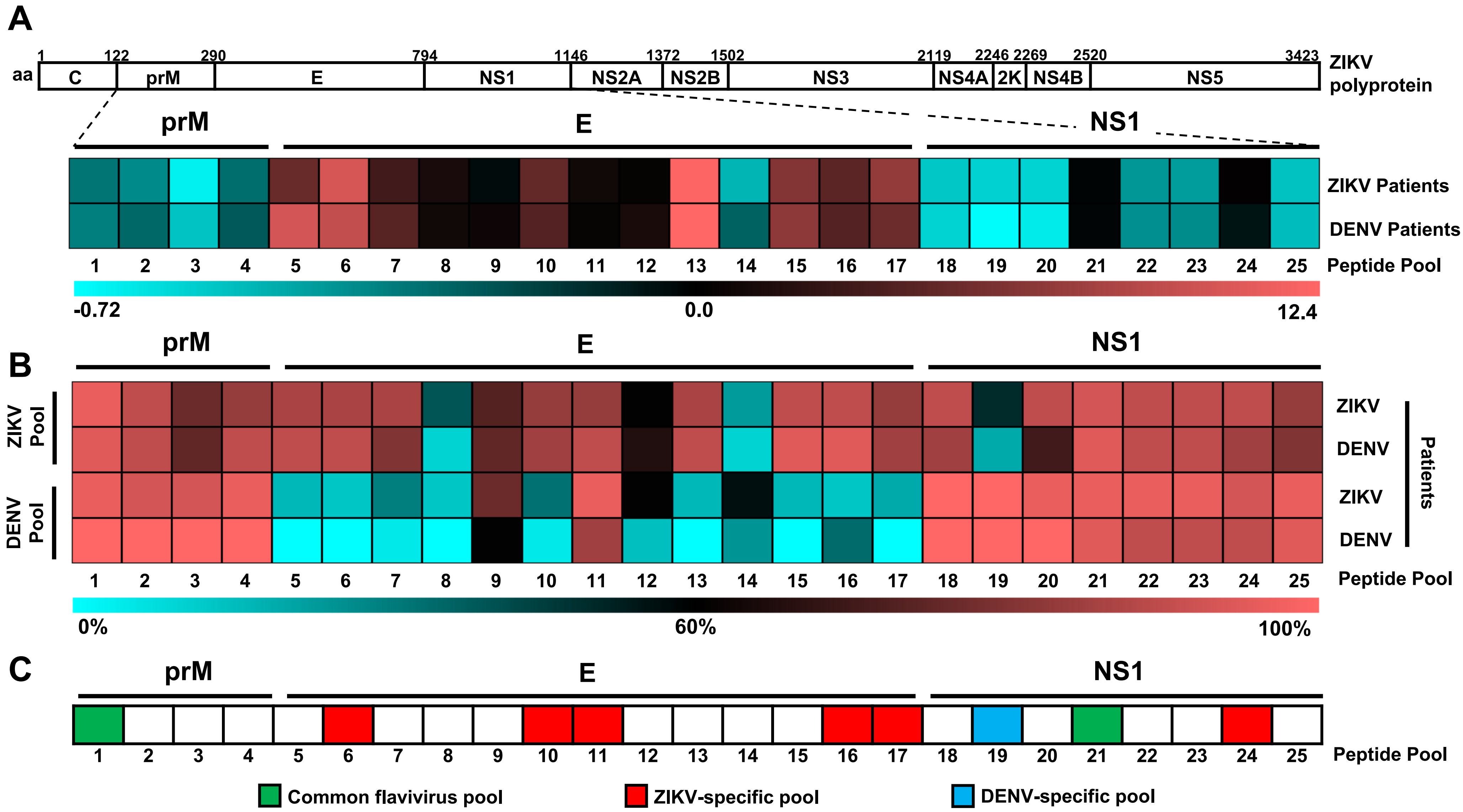


**Supplementary Figure 2. Preliminary mapping of common flavivirus, ZIKV-specific, and DENV-specific linear B-cell pooled epitopes within ZIKV and DENV proteome using ZIKV and DENV patient samples**. (A) Polyprotein of ZIKV H/PF/2013 (UniProtKB accession: A0A024B7W1), matched to results of pooled peptide-based ELISA experiments. Plasma samples of ZIKV patients (n=30) and serum samples of DENV patients (n=20) at 1:2000 dilution in duplicates were subjected to peptide-based ELISA using pooled peptides covering the precursor of membrane (prM; pools 1-4), envelope (E; pools 5-17) and non-structural 1 (NS1; pools 18-25) proteins of ZIKV and DENV proteome. Each pool consists of 5 peptides of 18-mer length, with overlapping sequence of 10 amino acids. IgG response of patients were normalized to mean of pooled healthy control. Patients' response to ZIKV and DENV pooled peptide-pairs were compared and the mean binding capacity are presented in a heat-map. A value of 0 on the scale denotes patients showing equal binding response to a ZIKV and DENV pooled peptide-pair, whereas values larger than 0 show preferential of patients to bind to ZIKV pooled peptide. Values smaller than 0 show binding preference of patients to DENV pooled peptide. (B) Percentage recognition of ZIKV and DENV patients to peptide pools were calculated. (C) A schematic representation to denote potential common flavivirus (in green), ZIKV-specific (in red), and DENV-specific pools (in blue) across prM, E and NS1 based on the heat-map analysis above.

**Supp. Figure 2 Amrun_Yee *et al.*, 2019**
