## Supplementary Figure 3 for "Novel differential linear B-cell epitopes to identify Zika and dengue virus infections in patients"

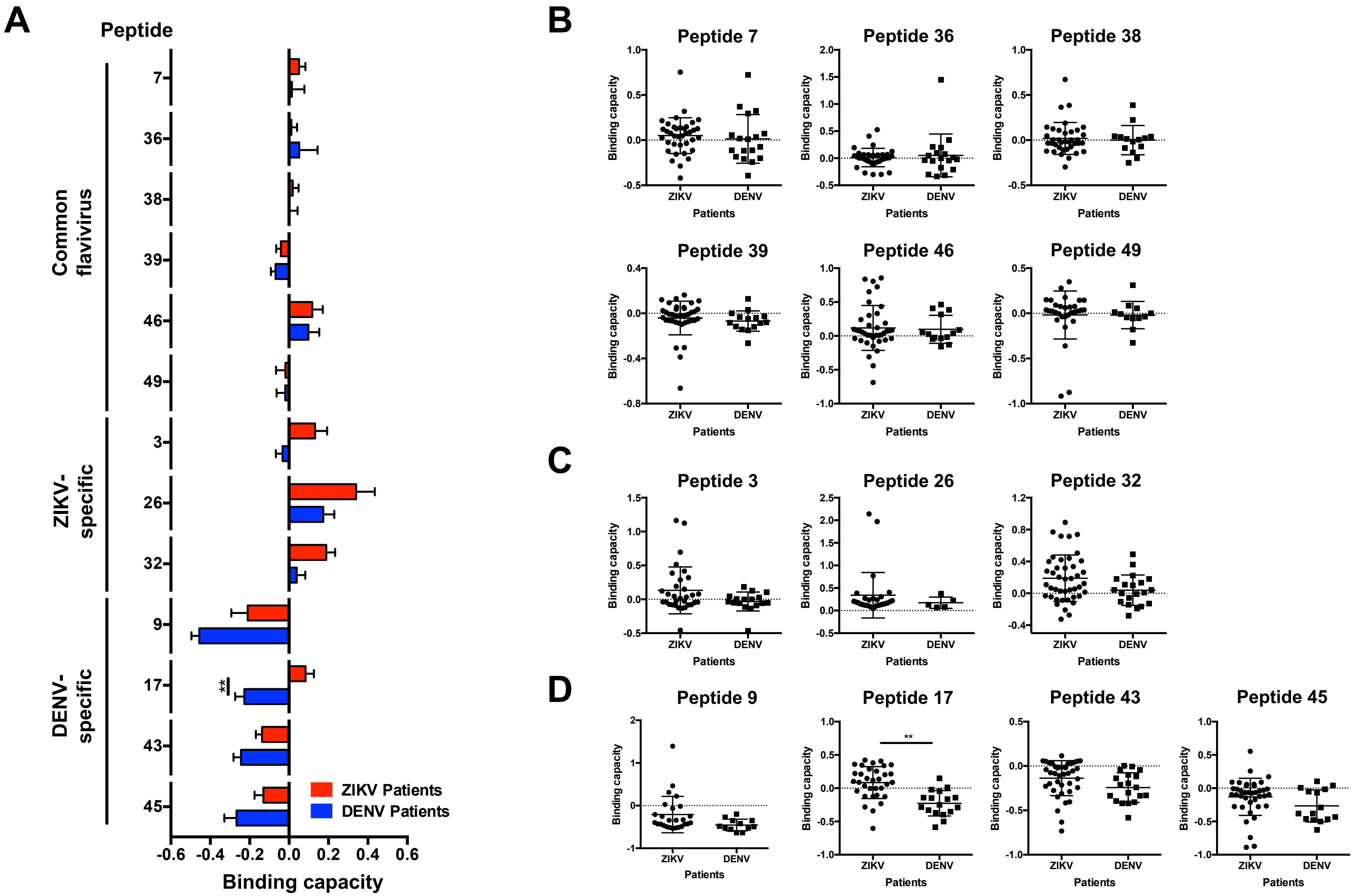


**Supplementary Figure 3. Peptide binding capacity of ZIKV and DENV patients on potential common flavivirus, ZIKV-specific, and DENV-specific linear B-cell epitopes.** Plasma samples of ZIKV (n=30-44) and serum samples of DENV (n=20) patients at late convalescent phase were tested at 1:2000 dilution in duplicates in a peptide-based ELISA, with pooled plasma of healthy donors used as negative control. The IgG binding capacity of patients positive for respective ZIKV and DENV peptide-pairs were calculated as [(ZIKV peptide response-DENV peptide response)/DENV peptide response] and the mean ± SEM values are presented in (A) for potential common flavivirus, ZIKV-specific, and DENV-specific linear B-cell epitopes. The distribution of binding capacity of individual ZIKV and DENV patients are shown in (B) for common flavivirus, (C) for ZIKV-specific and (D) for DENV-specific peptides. Data presented as mean ± SD. Statistical analysis was carried out using Mann-Whitney two-tailed test, with Bonferroni correction for multiple testing (***p*<0.01).

**Supp. Figure 3 Amrun_Yee *et al.*, 2019**
