## Supplementary Figure 4 for "Novel differential linear B-cell epitopes to identify Zika and dengue virus infections in patients"

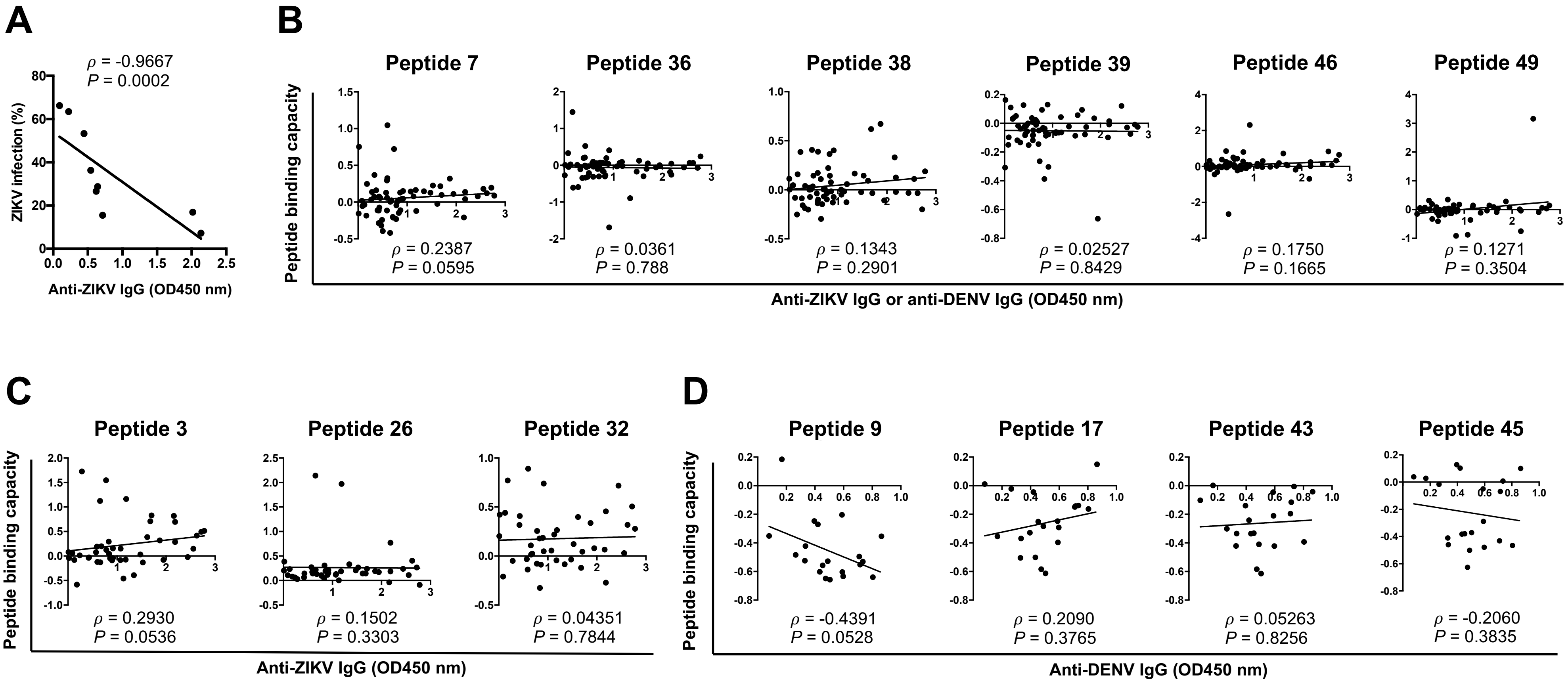


**Supplementary Figure 4. Correlation analysis of antibody and peptide response.** (A) Plasma samples of ZIKV patients (n=65) were pooled according to the levels of anti-ZIKV IgG titer (shown in Figure 1B). Mean neutralizing capacity of pooled ZIKV patients (shown in Figure 1E-F) and mean anti-ZIKV IgG levels of ZIKV patients were calculated for all time points (from acute to full recovery) and correlation was carried out. (B-D) Plasma samples from ZIKV (n=30-44) and serum samples from DENV (n=20) patients at late convalescent time point were tested for anti-ZIKV or anti-DENV IgG respectively at 1:2000 dilution, using purified ZIKV or DENV virions in a virion-based ELISA. In addition, plasma/serum samples were tested at 1:2000 dilution in duplicates for peptide-specific IgG using ZIKV and DENV peptides in a peptide-based ELISA. Patients' response to ZIKV and DENV peptide-pairs were compared and peptide binding capacity was calculated. Correlation of (B) both ZIKV and DENV patients' antibody response to common flavivirus peptides, (C) ZIKV patients' antibody response to ZIKV-specific peptides, and (D) DENV patients' antibody response to DENV-specific peptides. Correlation analysis was carried out using Spearman's rank correlation. Spearman's rho ($\rho$) and $p$-value ($P$) are presented.

**Supp. Figure 4 Amrun_Yee *et al.*, 2019**
