## Supplementary Table 1 for "Novel differential linear B-cell epitopes to identify Zika and dengue virus infections in patients"

### Supplementary Table 1: ZIKV and DENV peptides information

| Protein | Peptide no | Amino acid position on ZIKV (accession KJ776791) | | Corresponding ZIKV sequence | Corresponding DENV sequence | Peptide similarity (% identity, % query cover) | Classification of potential epitope |
|---|---|---|---|---|---|---|---|
| | | Start | End | | | | |
| prM | 1 | 6 | 23 | RGSAYYMYLDRNDAGEAI | RDGEPRMIVGKNERGKSL | No similarity | |
| | 2 | 15 | 32 | DRNDAGEAISFPTTLGMN | GKNERGKSLLFKTASGIN | No similarity | |
| | 3 | 24 | 41 | SFPTTLGMNKCYIQIMDL | LFKTASGINMCTLIAMDL | No similarity | **ZIKV-specific** |
| | 4 | 33 | 50 | KCYIQIMDLGHMCDATMS | MCTLIAMDLGEMCDDTVT | 80%, 55% | |
| | 5 | 42 | 59 | GHMCDATMSYECPMLDEG | GEMCDDTVTYKCPHITE | 62%, 72% | |
| | 6 | 51 | 68 | YECPMLDEGVEPDDVDCW | YKCPHITEVEPEDIDCW | 61%, 100% | |
| | 7 | 56 | 72 | LDEGVEPDDVDCWCNTT | HITEVEPEDIDCWCNLT | 82%, 64% | **Common flavivirus** |
| | 8 | 69 | 86 | CNTTSTWVVYGTCHHKKG | CNLTSTWVTYGTCNQAG | 73%, 83% | |
| | 9 | 78 | 92 | YGTCHHKKGEARRSR | TSTWVTYGTCNQAG | 67%, 40% | **DENV-specific** |
| | 10 | 100 | 118 | HSTRKLQTRSQTWLESREY | VGMGLDTRTQTWMSAEGAW | 67%, 47% | |
| E | 11 | 37 | 54 | DKPTVDIELVTTTVSNMA | NKPTLDIELQKTEATQLA | 78%, 50% | |
| | 12 | 61 | 72 | YEASISDMASDS | IEGKITNITTDS | No similarity | |
| | 13 | 73 | 90 | RCPTQGEAYLDKQSDTQY | RCPTQGEAVLPEEQDQNY | 100%, 44% | |
| | 14 | 91 | 108 | VCKRTLVDRGWGNGCGLF | VCKHTYVDRGWGNGCGLF | 89%, 100% | |
| | 15 | 113 | 130 | LVTCAKFACSKKMTGKSI | LVTCAKFQCLEPIEGKVV | 89%, 50% | |
| | 16 | 123 | 140 | KKMTGKSIQPENLEYRIM | EPIEGKVVQYENLKYTVI | 64%, 61% | |
| | 17 | 131 | 149 | PENLEYRIMLSVHGSQHS | YENLKYTVIITVHTGDQH | 53%, 88% | **DENV-specific** |
| | 18 | 149 | 166 | SGMIVNDTGHETDENRAK | GDQHQVGNETQGVTAEIT | No similarity | |
| | 19 | 157 | 174 | GHETDENRAKVEITPNSP | GNETQGVTAEITPQASTT | 100%, 22% | |
| | 20 | 166 | 183 | KVEITPNSPRAEATLGGF | TAEITPQASTTEAILPEY | 54%, 72% | |
| | 21 | 191 | 203 | EPRTGLDFSDLYY | SPRTGLDFNEMIL | 70%, 76% | |
| | 22 | 199 | 216 | SDLYYLTMNNKHWLVHKE | NEMILLTMKNKAWMVHRQ | 62%, 72% | |
| | 23 | 217 | 234 | WFHDIPLPWHAGADTGTP | WFFDLPLPWTSGATTETP | 67%, 100% | |
| | 24 | 235 | 245 | HWNNKEALVEF | TWNRKELLVTF | 70%, 90% | |
| | 25 | 244 | 261 | EFKDAHAKRQTVVVLGSQ | TFKNAHAKKQEVVVLGSQ | 82%, 94% | |
| | 26 | 271 | 288 | GALEAEMDGAKGRLSSGH | GATEIQNSGGTSIFAGH | 100%, 11% | **ZIKV-specific** |
| | 27 | 306 | 319 | SLCTAAFTFTKIPA | AMCTNTFVLKKEVS | No similarity | |
| | 28 | 325 | 342 | TVTVEVQYAGTDGPCKVP | TILIKVEYKGEDAPCKIP | 62%, 72% | |
| | 29 | 343 | 355 | AQMAVDMQTLTPV | FSTEDGQGKAHN | No similarity | |
| | 30 | 361 | 378 | ANPVITESTENSKMMLEL | ANPVVTKKEEPVNIEA | 83%, 33% | |
| | 31 | 402 | 419 | RSGSTIGKAFEATVRGAK | KKGSSIGKMFEATARGAR | 80%, 83% | |

| | | | | | | | |
|---|---|---|---|---|---|---|---|
| | 32 | 453 | 470 | FKSLFGGMSWFSQILIGT | YTALFSGVSWVMKIGIGV | 71%, 38% | **ZIKV-specific** |
| NS1 | 33 | 1 | 18 | DVGCSVDFSKKETRCGTG | DMGCVINWKGKELKCGSG | 44%, 100% | |
| | 34 | 19 | 36 | VFVYNDVEAWRDRYKYHP | IFVTNEVHTWTEQYKFQA | 41%, 94% | |
| | 35 | 55 | 72 | CGISSVSRMENIMWRSVE | CGIRSTTRMENLLWKQIA | 64%, 77% | |
| | 36 | 70 | 85 | SVEGELNAILEENGVQ | QIANELNYILWENNIK | 70%, 56% | **Common flavivirus** |
| | 37 | 91 | 112 | GSVKNPMWRGPQRLPVPVNELP | GDIIGVLEQGKRTLTPQPMELK | No similarity | |
| | 38 | 119 | 136 | GKSYFVRAAKTNNSFVVD | GKAKIVTAETQNSSFIID | 57%, 38% | **Common flavivirus** |
| | 39 | 137 | 154 | GDTLKECPLKHRAWNSFL | GPNTPECPSASRAWNVWE | 70%, 55% | **Common flavivirus** |
| | 40 | 155 | 176 | VEDHGFGVFHTSVWLKVREDYS | VEDYGFGVFTTNIWLKLREVYT | 71%, 95% | |
| | 41 | 239 | 256 | SDLIIPKSLAGPLSHHNT | SDMIIPKSLAGPISQHNH | 82%, 94% | |
| | 42 | 248 | 265 | AGPLSHHNTREGYRTQMK | AGPISQHNHRPGYHTQTA | 69%, 88% | |
| | 43 | 257 | 274 | REGYRTQMKGPWHSEELE | RPGYHTQTAGPWHLGKLE | 69%, 72% | **DENV-specific** |
| | 44 | 270 | 281 | SEELEIRFEECP | LGKLELDFNYCE | No similarity | |
| | 45 | 275 | 292 | IRFEECPGTKVHVEETCG | LDFNYCEGTTVVITENCG | 54%, 72% | **DENV-specific** |
| | 46 | 284 | 301 | KVHVEETCGTRGPSLRST | TVVITENCGTRGPSLRTT | 85%, 72% | **Common flavivirus** |
| | 47 | 293 | 310 | TRGPSLRSTTASGRVIEE | TRGPSLRTTTVSGKLIHE | 72%, 100% | |
| | 48 | 302 | 319 | TASGRVIEEWCCRECTMP | TVSGKLIHEWCCRSCTLP | 67%, 100% | |
| | 49 | 315 | 326 | ECTMPPLSFRAK | SCTLPPLRYMGE | 83%, 50% | **Common flavivirus** |
| | 50 | 320 | 337 | PLSFRAKDGCWYGMEIRP | PLRYMGEDGCWYGMEIRP | 72%, 100% | |
| | 51 | 338 | 353 | RKEPESNLVRSMVTAG | ISEKEENMVKSLVSAG | No similarity | |
