## Supplementary Table 2 for "Novel differential linear B-cell epitopes to identify Zika and dengue virus infections in patients"

**Supplementary Table 2: Preliminary results of ZIKV and DENV patients' response to ZIKV and DENV pooled peptides**

| Protein | Peptide Pool | Percentage recognition (%)[a] | | | | Mean binding capacity[b] | | Relative difference[c] | Preliminary classification of peptide pool[d] |
|---|---|---|---|---|---|---|---|---|---|
| | | ZIKV Patients (n=30) | | DENV Patients (n=20) | | | | | |
| | | ZIKV Pool | DENV Pool | ZIKV Pool | DENV Pool | ZIKV Patients | DENV Patients | | |
| prM | 1 | 96.7 | 96.7 | 95 | 100 | -0.325 | -0.360 | 0.035 | **Common** |
| | 2 | 90.0 | 93.3 | 90 | 100 | -0.392 | -0.295 | 0.097 | |
| | 3 | 76.7 | 93.3 | 75 | 100 | -0.684 | -0.553 | 0.131 | |
| | 4 | 83.3 | 96.7 | 90 | 100 | -0.313 | -0.254 | 0.059 | |
| E | 5 | 86.7 | 16.7 | 90 | 0 | 5.066 | 10.343 | 5.277 | |
| | 6 | 86.7 | 13.3 | 90 | 0 | 10.379 | 9.512 | 0.867 | **ZIKV-specific** |
| | 7 | 86.7 | 30.0 | 80 | 5 | 3.102 | 4.222 | 1.120 | |
| | 8 | 40.0 | 13.3 | 10 | 0 | 1.212 | 0.741 | 0.471 | |
| | 9 | 73.3 | 76.7 | 75 | 60 | -0.029 | 0.653 | 0.681 | |
| | 10 | 83.3 | 33.3 | 85 | 5 | 4.730 | 4.043 | 0.687 | **ZIKV-specific** |
| | 11 | 83.3 | 96.7 | 90 | 85 | 0.763 | 0.301 | 0.462 | **ZIKV-specific** |
| | 12 | 60.0 | 60.0 | 65 | 15 | 0.267 | 1.222 | 0.954 | |
| | 13 | 86.7 | 16.7 | 90 | 0 | 12.259 | 12.399 | 0.140 | |
| | 14 | 23.3 | 56.7 | 10 | 25 | -0.513 | -0.279 | 0.234 | |
| | 15 | 90.0 | 16.7 | 95 | 0 | 6.344 | 6.814 | 0.470 | |
| | 16 | 90.0 | 13.3 | 95 | 35 | 4.429 | 4.040 | 0.388 | **ZIKV-specific** |
| | 17 | 83.3 | 20.0 | 85 | 0 | 7.079 | 5.210 | 1.869 | **ZIKV-specific** |
| NS1 | 18 | 90.0 | 100.0 | 85 | 100 | -0.562 | -0.601 | 0.039 | |
| | 19 | 50.0 | 100.0 | 20 | 100 | -0.595 | -0.716 | 0.120 | **DENV-specific** |
| | 20 | 90.0 | 96.7 | 70 | 100 | -0.600 | -0.665 | 0.065 | |
| | 21 | 93.3 | 96.7 | 95 | 95 | -0.014 | -0.016 | 0.002 | **Common** |
| | 22 | 90.0 | 96.7 | 90 | 90 | -0.427 | -0.404 | 0.022 | |
| | 23 | 90.0 | 96.7 | 90 | 90 | -0.445 | -0.396 | 0.049 | |
| | 24 | 90.0 | 93.3 | 85 | 90 | 0.134 | -0.055 | 0.189 | **ZIKV-specific** |
| | 25 | 83.3 | 96.7 | 80 | 95 | -0.548 | -0.533 | 0.015 | |

[a]Patient samples are positive if their normalized peptide responses (calculated as OD of patient sample/mean OD of pooled healthy) are more than 1.01.

[b]Binding capacity of a patient positive for a peptide-pair was calculated as: normalized values of [(ZIKV peptide response-DENV peptide response)/DENV peptide response].
Values close to 0 denote equal binding of patient to ZIKV and DENV peptide.
Values more than 0 denote a patient's preference to bind to ZIKV peptide more than DENV peptide.
Values less than 0 denote a patient's preference to bind to DENV peptide more than ZIKV peptide.

[c]Relative difference is calculated as the diference in the mean binding capacity of ZIKV patients and DENV patients. Values are rounded up to 3 decimal places.

[d]Common flavivirus epitopes: ≥ 60% of ZIKV and DENV patients recognize both ZIKV and DENV peptides of peptide-pair; ZIKV-specific epitopes: ≥ 60% of ZIKV patients recognize at least ZIKV peptide of peptide-pair; DENV-specific epitopes: ≥ 60% of DENV patients recognize at least DENV peptide of peptide-pair.
