## Supplementary Table 3 for "Novel differential linear B-cell epitopes to identify Zika and dengue virus infections in patients"

**Supplementary Table 3: Evaluation of patients of different diagnoses and cohorts with potential linear B-cell epitopes**

| Cohort | Patient | Peptide | | | | | | | | | | | | |
|---|---|---|---|---|---|---|---|---|---|---|---|---|---|---|
| | | Common flavivirus[a] | | | | | | ZIKV-specific[b] | | | DENV-specific[b] | | | |
| | | 7 | 36 | 38 | 39 | 46 | 49 | 3 | 26 | 32 | 9 | 17 | 43 | 45 |
| ZIKV (Singapore) | 1 | y | y | y | y | y | y | e | z | e | d | z | e | e |
| | 2 | y | y | y | y | n | n | z | n | z | d | d | d | d |
| | 3 | y | y | y | y | y | n | z | z | d | n | z | d | d |
| | 4 | y | y | y | y | y | y | z | z | z | n | z | e | d |
| | 5 | y | y | y | y | y | y | e | z | e | d | z | d | d |
| | 6 | y | y | y | y | y | y | z | z | z | n | e | e | d |
| | 7 | y | y | y | y | y | y | z | z | z | n | e | e | e |
| | 8 | y | n | y | n | y | y | n | n | z | n | d | d | n |
| | 9 | y | y | y | y | y | y | z | z | e | e | z | d | z |
| | 10 | y | y | y | y | y | y | e | z | e | d | z | d | e |
| DENV (Singapore) | 1 | y | y | y | y | y | y | e | e | z | d | z | e | z |
| | 2 | y | n | n | n | n | n | d | n | z | n | n | n | n |
| | 3 | y | y | y | n | n | n | z | n | z | d | d | d | d |
| | 4 | y | y | y | y | y | y | e | n | e | d | d | d | d |
| | 5 | y | y | y | y | y | y | e | z | e | d | d | d | d |
| | 6 | y | n | n | n | n | n | e | n | z | n | n | d | d |
| | 7 | y | y | y | n | y | y | e | n | d | d | e | d | n |
| | 8 | y | n | y | y | y | n | z | n | d | n | d | e | n |
| | 9 | y | y | y | y | y | y | e | z | d | d | d | e | e |
| | 10 | y | y | y | y | y | y | d | e | d | d | e | d | e |
| DENV (Thailand) | 1 | n | y | n | n | n | n | n | n | d | n | n | d | n |
| | 2 | y | y | n | y | n | n | z | n | d | z | d | d | z |
| | 3 | y | n | n | y | n | n | z | n | d | d | d | d | n |
| | 4 | n | n | n | n | n | n | n | n | e | z | d | d | n |
| | 5 | n | n | n | n | n | n | e | n | e | z | d | d | n |
| Bacteria (Thailand) | 1 | y | n | n | y | n | n | d | n | d | z | d | d | n |
| | 2 | n | n | n | n | n | n | d | n | d | z | d | d | n |
| | 3 | y | n | n | n | n | n | e | n | d | z | d | d | n |
| | 4 | y | y | y | y | y | y | e | n | d | z | d | d | e |
| | 5 | y | n | n | n | n | n | d | n | d | z | d | d | n |
| Unknown (Thailand) | 1 | y | y | y | y | y | n | d | z | d | d | d | z | d |
| | 2 | n | n | n | n | n | n | n | n | z | n | d | d | n |
| | 3 | y | y | y | y | y | n | d | n | d | d | d | d | n |
| | 4 | y | n | y | y | n | n | d | n | d | z | d | d | n |
| | 5 | y | n | y | y | n | n | n | n | d | n | d | d | n |
| | 6 | y | y | y | y | y | y | e | z | d | d | d | d | e |
| | 7 | n | n | n | n | n | n | n | n | d | n | n | n | n |
| | 8 | n | n | n | n | y | n | d | n | d | n | n | d | n |

[a]A patient sample is considered positive (indicated as "y") if it has a normalized peptide response higher than pooled healthy control for both ZIKV and DENV peptide-pair. If a sample peptide-pair response is lower than the healthy, it is considered negative (indicated as "n").

[b]If a patient sample is positive for a peptide, i.e. has a higher normalized peptide response than pooled healthy control, the binding capacity of peptides (calculated as [(ZIKV peptide response-DENV peptide response)/DENV peptide response] was then determined. For patients with peptide binding capacity values of
i) binding capactiy ≥ 0.1 → positive for ZIKV peptide (indicated as "z")
ii) binding capacity ≤ 0.1 → positive for DENV peptide (indicated as "d")
ii) -0.1 < binding capacity < 0.1 → equal recognition of ZIKV and DENV peptide-pair (indicated as "e")
