## Supplementary Table 4 for "Novel differential linear B-cell epitopes to identify Zika and dengue virus infections in patients"

**Supplementary Table 4: Evaluation of common and differential flavivirus peptide mix with patient cohorts**

| Cohort | Patient | Percentage of positive peptides (%) | | | Outcome of patients | | | |
|---|---|---|---|---|---|---|---|---|
| | | Common flavivirus[a] | ZIKV-specific[b] | DENV-specific[c] | Common flavivirus[a] | ZIKV-specific[b] | DENV-specific[c] | Final "diagnosis"[d] |
| ZIKV (Singapore) | 1 | 100 | 33.3 | 100 | y | n | y | d |
| | 2 | 50 | 66.7 | 100 | n | y | y | n |
| | 3 | 100 | 66.7 | 0 | y | y | n | z |
| | 4 | 100 | 100.0 | 0 | y | y | n | z |
| | 5 | 100 | 33.3 | 100 | y | n | y | d |
| | 6 | 100 | 100.0 | 0 | y | y | n | z |
| | 7 | 100 | 100.0 | 0 | y | y | n | z |
| | 8 | 75 | 33.3 | 0 | y | n | n | n |
| | 9 | 100 | 66.7 | 0 | y | y | n | z |
| | 10 | 100 | 33.3 | 100 | y | n | y | d |
| DENV (Singapore) | 1 | 100 | 33.3 | 100 | y | n | y | d |
| | 2 | 0 | 33.3 | 0 | n | n | n | n |
| | 3 | 50 | 66.7 | 100 | n | y | y | n |
| | 4 | 100 | 0.0 | 100 | y | n | y | d |
| | 5 | 100 | 33.3 | 100 | y | n | y | d |
| | 6 | 25 | 33.3 | 0 | n | n | n | n |
| | 7 | 75 | 0.0 | 100 | y | n | y | d |
| | 8 | 75 | 33.3 | 0 | y | n | n | n |
| | 9 | 100 | 33.3 | 100 | y | n | y | d |
| | 10 | 100 | 0.0 | 100 | y | n | y | d |
| DENV (Thailand) | 1 | 25 | 0.0 | 0 | n | n | n | n |
| | 2 | 75 | 33.3 | 0 | y | n | n | n |
| | 3 | 50 | 33.3 | 100 | n | n | y | n |
| | 4 | 25 | 0.0 | 0 | n | n | n | n |
| | 5 | 25 | 0.0 | 0 | n | n | n | n |
| Bacteria (Thailand) | 1 | 50 | 0.0 | 0 | n | n | n | n |
| | 2 | 25 | 0.0 | 0 | n | n | n | n |
| | 3 | 25 | 0.0 | 0 | n | n | n | n |
| | 4 | 100 | 0.0 | 0 | y | n | n | n |
| | 5 | 25 | 0.0 | 0 | n | n | n | n |
| Unknown (Thailand) | 1 | 100 | 33.3 | 100 | y | n | y | d |
| | 2 | 25 | 33.3 | 0 | n | n | n | n |
| | 3 | 100 | 0.0 | 100 | y | n | y | d |
| | 4 | 75 | 0.0 | 0 | y | n | n | n |
| | 5 | 50 | 0.0 | 0 | n | n | n | n |
| | 6 | 100 | 33.3 | 100 | y | n | y | d |
| | 7 | 25 | 0.0 | 0 | n | n | n | n |
| | 8 | 50 | 0.0 | 0 | n | n | n | n |

Based on Supplementary Table 3, the percentage of positive peptides were calculated and "outcome" of patients were assigned.

[a]If sample is positive for 3 or more (out of 4) common flavivirus peptides, i.e. ≥75% positive, the patient is considered positive (indicated as "y" in the outcome) for flavivirus infection. If sample is ≤75% positive, the patient is considered negative (indicated as "n" in the outcome).

[b]If sample is positive for 2 or more (out of 3) ZIKV-specific peptides, i.e. ≥66.7% positive, the patient is considered positive (indicated as "y" in the outcome) for ZIKV infection. If sample is ≤66.7% positive, the patient is considered negative (indicated as "n" in the outcome).

[c]If sample is positive for the DENV-specific peptide, i.e. ≥100% positive, the patient is considered positive (indicated as "y" in the outcome) for DENV infection, whereas negative for the DENV-specific peptide is indicated as "n" in the outcome.

[d]Based on the respective outcomes of the epitope categories, the following combinations produce the final ZIKV and DENV "diagnosis" of patients.

<table>
<tr><th>Common flavivirus</th><th>ZIKV-specific</th><th>DENV-specific</th><th>Final "diagnosis"</th></tr>
<tr><td>y</td><td>y</td><td>n</td><td>ZIKV positive ("z")</td></tr>
<tr><td>y</td><td>n</td><td>y</td><td>DENV positive ("d")</td></tr>
<tr><td>y</td><td>n</td><td>n</td><td rowspan="4">ZIKV and DENV negative ("n")</td></tr>
<tr><td>n</td><td>y</td><td>n</td></tr>
<tr><td>n</td><td>n</td><td>y</td></tr>
<tr><td>n</td><td>n</td><td>n</td></tr>
</table>
